# CD3⁺CD4⁻CD8⁻ (Double-Negative) T-cells Provide Inflammatory Signals That Influence Myeloid Cells and Enhance Survival in Glioblastoma

**DOI:** 10.64898/2026.11.16.767095

**Authors:** Malcolm F. McDonald, Rajan Patel, Jack Tremblay, Khatri Latha, Yuhui Yang, Ashley Puentes, Arya Shetty, Yi-Cian Zheng, Victoria Soueng, Rouchen Du, Harrshavasan Congivaram, Hinda Najem, Eric A. Goethe, Rebecca Murdagh, Yeunjung Ko, Akdes Serin Harmanci, Benjamin Deneen, Jamie Anastas, Amy B. Heimberger, Ganesh Rao

## Abstract

Glioblastoma (GBM) remains resistant to immunotherapy, in part because of a profoundly immunosuppressive tumor microenvironment. While investigating the consequences of conventional T-cell depletion in a murine glioma model, we unexpectedly identified a population of double-negative T-cells (DNTs; CD3⁺CD4⁻CD8⁻) that persisted following combined depletion of CD4⁺ and CD8⁺ T-cells. DNT enrichment was associated with significantly prolonged survival and revealed a distinct inflammatory transcriptional program characterized by activation-associated gene expression. Single-cell transcriptomic analyses demonstrated extensive remodeling of the glioma myeloid compartment, including enrichment of inflammatory signaling pathways and reduced expression of immunosuppressive markers. In complementary human studies, unmodified healthy donor-derived DNTs suppressed glioma stem cell viability, enhanced macrophage phagocytic activity, and reduced expression of the inhibitory myeloid receptor LAIR1. Intracranial administration of these unmodified DNTs significantly prolonged survival compared with PanT cells in an orthotopic glioblastoma xenograft model.. Collectively, these findings identify DNTs as previously underappreciated regulators of glioblastoma immunity that exert antitumor effects through both direct tumor suppression and inflammatory remodeling of the glioma myeloid compartment. These results support further investigation of DNT-based cellular therapies for the treatment of glioblastoma.

## Introduction

Glioblastoma (GBM) is the most common and aggressive primary malignant brain tumor in adults^1^. Despite maximal safe surgical resection followed by radiation and temozolomide chemotherapy, median survival remains approximately 15–20 months, and long-term survival is rare^2^. Efforts to improve outcomes through immunotherapy have achieved limited success, largely due to the profoundly immunosuppressive nature of the glioblastoma microenvironment^3–6^. A defining feature of the glioblastoma microenvironment is the abundance of immunosuppressive myeloid cells, including tumor-associated macrophages and microglia, which can comprise up to half of the tumor mass^7–9^. These populations promote tumor growth through suppression of effector immune responses, production of immunoregulatory cytokines, and support of tumor invasion and angiogenesis^6,10–14^. Consequently, increasing attention has focused on understanding the interactions between lymphocytes and myeloid cells within gliomas and identifying immune populations capable of overcoming this suppressive microenvironment.

Our laboratory previously demonstrated that genetic deletion of CD8⁺ T-cells significantly altered the glioma myeloid compartment, resulting in increased expression of inflammatory markers including MHC II and TNFα^15^. Surprisingly, however, these changes failed to improve survival, suggesting that conventional T-cell populations exert complex and potentially opposing effects on glioma progression. These findings raised the possibility that additional lymphocyte populations may contribute to antitumor immunity and myeloid regulation within glioblastoma. To further investigate the contribution of conventional T-cell subsets to glioma immunity, we depleted CD4⁺ T-cells (ΔCD4) in CD8 knockout mice (CD8^-/-^) bearing RCAS/tv-a PDGFB/STAT3-driven gliomas with the goal of evaluating tumor progression following near-complete elimination of conventional T-cells.

Although combined depletion of CD4⁺ and CD8⁺ T-cells increased overall survival, it did not eliminate intra-tumoral T lymphocytes. Instead, we identified a persistent population of CD3⁺ T-cells that lacked expression of both CD4 and CD8. Phenotypic characterization revealed that these cells were consistent with double-negative T-cells (DNTs), a rare but increasingly recognized T-cell subset that has been implicated in immune regulation, transplantation tolerance, autoimmunity, and antitumor immunity^16–25^. However, the role of DNTs in glioblastoma and their interactions with the glioma immune microenvironment remain poorly understood.

Here, we show that CD4 depletion in CD8-deficient glioma-bearing mice unexpectedly prolongs survival and reveals a persisting inflammatory DNT population. DNT enrichment is accompanied by substantial changes in the glioma myeloid compartment. In complementary human experiments, unmodified healthy donor-derived DNTs suppress glioma stem cell viability and alter macrophage phenotype and phagocytic activity. These cells also prolong survival in an orthotopic glioblastoma xenograft model without CAR engineering. Together, these findings connect the biology of DNTs within gliomas to the therapeutic potential of unmodified donor-derived DNTs.

## Results

### Double-negative T-cell enrichment is associated with improved survival in glioblastoma

Dysfunctional T-cells are known to play an important role in the immunosuppressive environment; however, previous work in the lab established that knockout of CD8 (CD8^-/-^) provided no survival benefit in a syngeneic model of high-grade glioma although an increase in inflammatory TAMs was observed. To further dissect the role of T-cells in GBM survival, we removed CD4^+^ and CD8^+^ T-cells from the tumor microenvironment using a combined CD8^- /-^ germline knockout and treatment with an anti-CD4 antibody (ΔCD4) in a syngeneic, immunocompetent RCAS model of high-grade glioma (Figure 1a). Dual overexpression of PDGFβ and STAT3 in mice expressing TVA under the nestin promoter (nTVA) faithfully recapitulates histopathological signatures of high-grade glioma in an immunocompetent model^26^. Mice received ΔCD4 or isotype control IgG antibodies treatment every 5 days starting at P10 for 3 weeks (Figure 1a). ΔCD4 treatment successfully removed CD4^+^ T-cells from the microenvironment (Supp Figure 1a-b). Unexpectedly, combined CD4 depletion in CD8^-/-^ mice produced a marked survival advantage in the ΔCD4/CD8^-/-^ mice compared to both IgG/CD8^+/+^ (p = 0.0256, Log-rank) and ΔCD4/CD8^+/+^ mice (p = 0.0008, Log-rank) (Figure 1b). CD4 depletion alone had no survival benefit. Furthermore, in human IDH WT GBM, combined expression of CD4, CD8A, and CD8B stratified survival with the bottom quartile of expression providing survival benefit over the top quartile (p = 0.011, Log-rank) (Figure 1c). Removal of CD4^+^ and CD8^+^ T-cells created a survival benefit in a syngeneic, immunocompetent murine glioma model. Collectively, these findings suggest that removal of conventional CD4^+^ and CD8^+^ T-cell populations unexpectedly enhances survival in glioma-bearing mice and suggests that alternative immune populations may contribute to antitumor immunity in glioblastoma. These observations prompted further investigation into the immune composition of ΔCd4/Cd8^−/−^ tumors.

**Figure 1:**
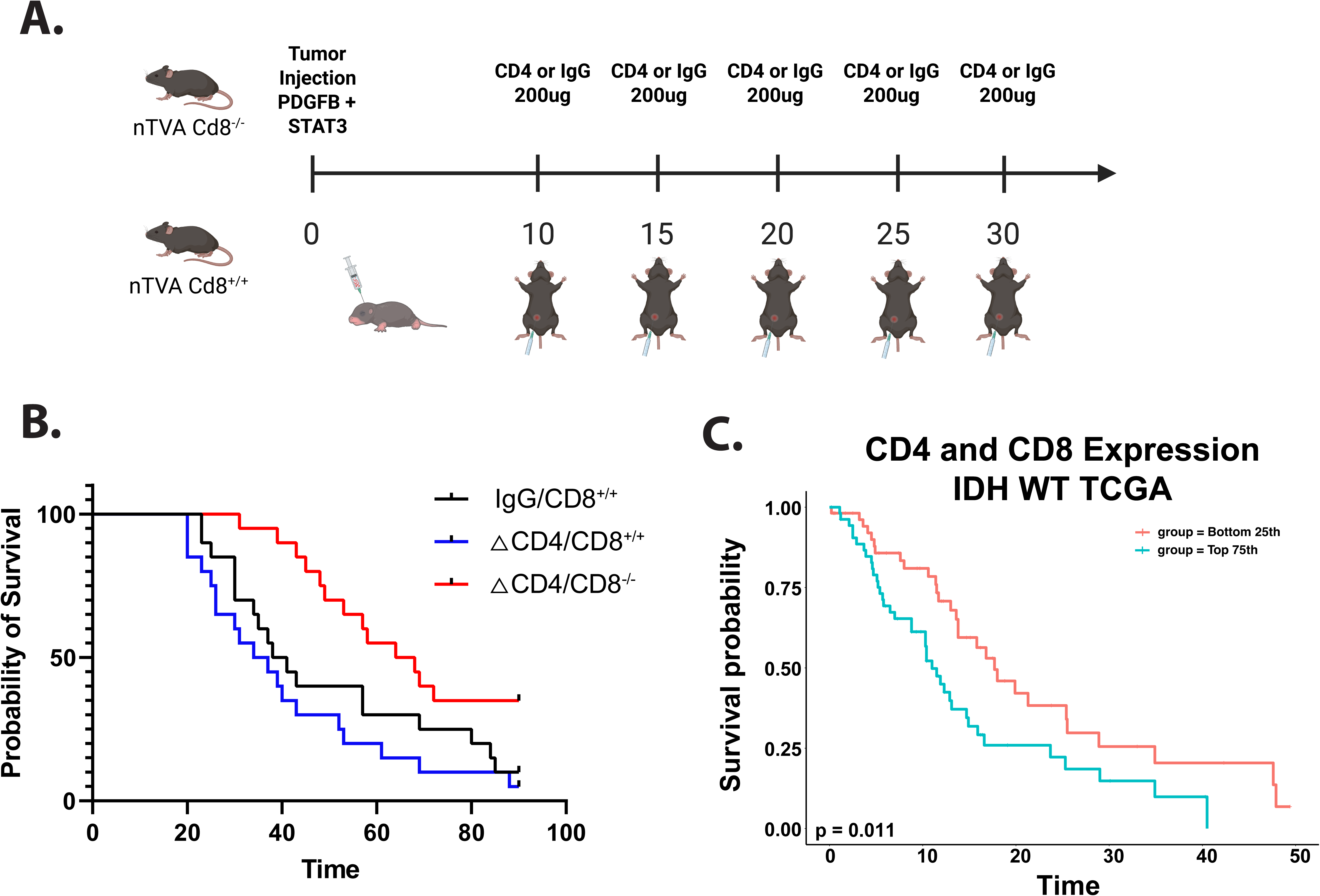
DNT enrichment is associated with improved survival. (A). Schematic for depletion of T-cells from RCAS model of gliomagenesis. (B). Improved survival for ΔCD4/Cd8^-/-^(med OS = 66 days) compared to IgG/Cd8^+/+^ (med OS = 39.5 days, p=0.0256, log-rank) and ΔCD4/Cd8^+/+^(med OS = 35.5 days, p=0.0008, log-rank) but no difference between ΔCD4/Cd8^+/+^ and IgG/Cd8^+/+^ (p=0.2892, log-rank). (C). Top quartile expression for combined single sample GSEA scoring for CD4, CD8A, and CD8B serves as a negative prognostic indicator for survival in TCGA IDH WT GBM compared to bottom quartile of expression (p=0,011, log-rank).

### DNTs exhibit an activated inflammatory transcriptional program distinct from conventional tumor-infiltrating lymphocytes

To investigate specific changes in the immune microenvironment induced by CD4^+^ and CD8^+^ T-cell removal, we conducted single cell RNA sequencing (scRNAseq) of the intratumoral immune population using a Percoll enriched gradient from tumor bearing mice at P45 (Figure 2a-b). This timepoint captured a snapshot of the microenvironment as the survival curves begin to separate. We were able to detect many immune constituents including robust representations of tumor associated macrophages and microglia as well as tumor infiltrating lymphocytes (TILs) (Figure 2b). Despite the removal of CD4^+^ and CD8^+^ T-cells, we detected tumor-infiltrating CD3^+^ T-cells (Figure 2c). These CD3^+^ T-cells, double negative for CD4^+^ and CD8^+^ (DNTs), lacked markers consistent with NK-cells, NKT-cells, or γδ T-cells (Figure 2d, Supp Figure 2a).

**Figure 2:**
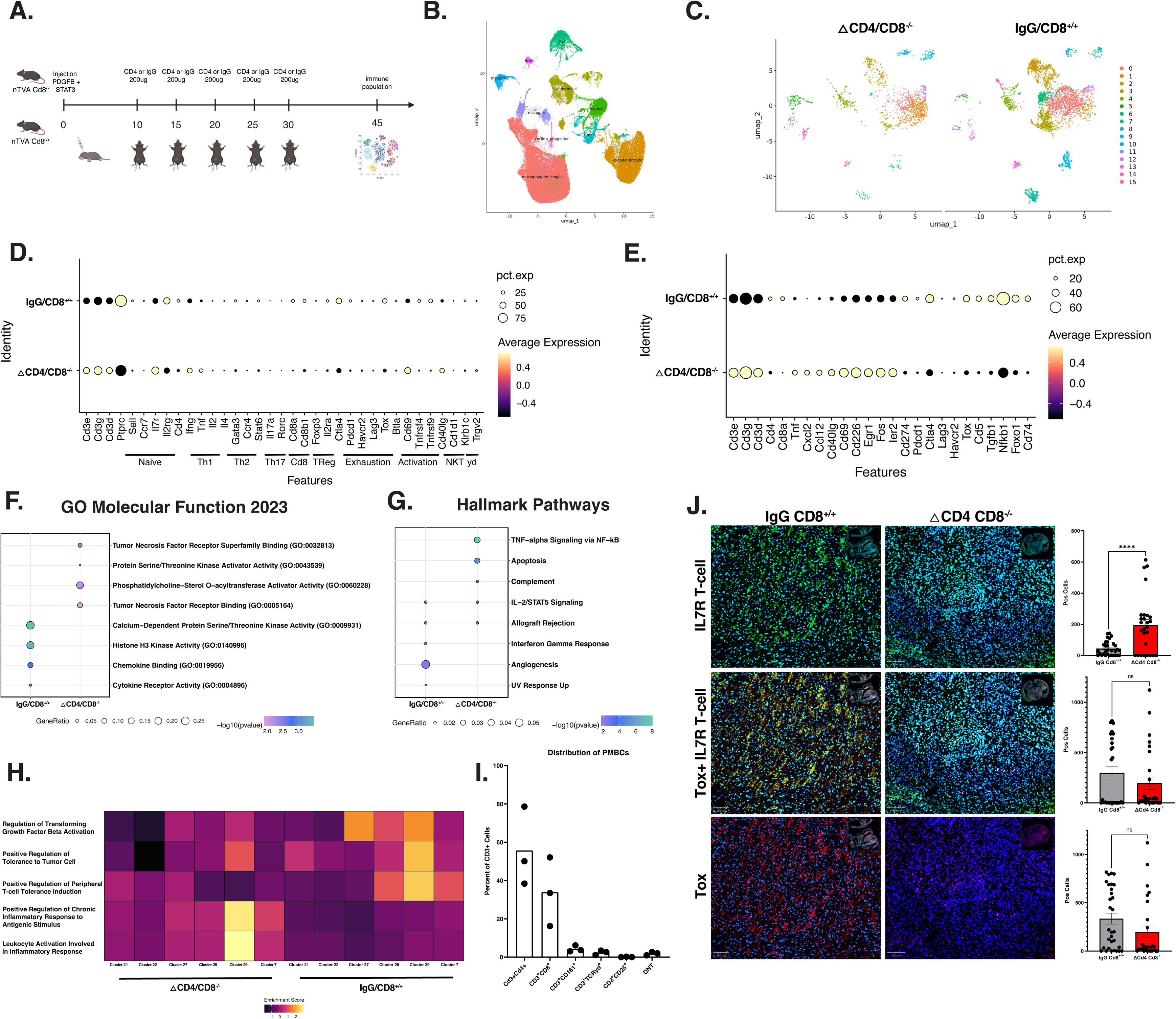
scRNAseq reveals population of inflammatory double negative T-cells. (A). Schematic for depletion scRNAseq from T-cell depleted mice. (B). Major cell types of the Seurat clusters. (C). Transcriptional differences between ΔCD4/Cd8^-/-^ and IgG/Cd8^+/+^ T-cells (D). Expression of major T-cell subtype markers in T-cells between ΔCD4/Cd8^-/-^ and IgG/Cd8^+/+^. (E). Differentially expressed genes (DEGs) between ΔCD4/Cd8^-/-^ and IgG/Cd8^+/+^. (F). Gene ontologies for DEGs from each group for Molecular Function and Hallmark pathways. (H). Single cell Gene Set Enrichment Analysis for different T-cell clusters by experiment. (I). Quantification of flow cytometry of isolated peripheral derived mononuclear cells (PBMC) from glioblastoma patients. (J). Multiplex fluorescence imaging for intratumoral IL7R T-cells and Tox+ T-cells demonstrating increased tumor infiltrating non-exhausted T-cells in ΔCD4/Cd8^- /-^ compared to IgG/Cd8^+/+^ (p-val < 0.0001, t-test) with relatively similar number of Tox+ T-cells (p-val = 0.2474, t-test).

To further characterize the potential function of these cells, we investigated their transcriptional patterns. Unlike the signatures of well-established tumor infiltrating CD3^+^ T-cells in glioma, DNTs demonstrated robust upregulation of pro-inflammatory markers (Tnf, Ccl12) and T-cell activation markers (Cd40lg, Il7r, Cd69, Cd226, Egr1, Fos, Ler2) (Figure 2e, Supp Figure 2b). Furthermore, DNTs demonstrated enrichment of TNFα and complement signaling programs in Molecular Function Ontologies and Hallmark Pathways (Figure 2f-g). Specific DNT clusters highly upregulated inflammatory and activating leukocyte signaling patterns in single-cell gene set enrichment analysis (scGSEA), demonstrating robust upregulation of inflammatory signaling and leukocyte activation (Figure 2h). DNTs have transcriptional patterns consistent with activated T-cells, with robust expression of inflammatory programs and cytokine expression.

Compared to DNTs, the conventional TILs from IgG/CD8^+/+^ demonstrated characteristic patterns of T-cell suppression and exhaustion outlined in other studies. IgG/CD8^+/+^ upregulated markers of T-cell exhaustion (Pdcd1, Ctla4, Tox) and immunosuppression (Tgfb1, Nfkb1, Cd5) (Figure 2e). The IgG/CD8^+/+^ TILs expressed higher signaling programs for chemokine binding and tumor-supportive activity related to angiogenesis (Figure 2f-g). In contrast to the inflammatory DNT clusters in scGSEA, IgG/Cd8^+/+^ T-cells upregulated pathways related to TGFβ activity and tolerance to tumor cells (Figure 2h). These different transcriptional signatures appeared to create a dichotomy between tumor suppressive activity in DNTs and tumor supportive activity in the IgG Cd8^+/+^ TILs that could influence the observed difference in survival.

We sought to understand if these cells exist in human patients. FACS analysis of peripheral blood from glioma patients demonstrated that a similar population of DNT exists in at about 2% of CD3^+^ cells (Figure 2i, Supp Figure 2c). While these cells were rare in patients, we wanted to understand the spatial representation of these T-cells within tumors. Utilizing multiplex immunofluorescent imaging in our mouse models, we found that there was an increase in the number of non-exhausted IL7R^+^ T-cells within ΔCD4/Cd8^-/-^tumors compared IgG/Cd8^+/+^, while both displayed relatively similar levels of exhausted Tox^+^IL7R^+^ T-cells (Figure 2j). DNTs appear to be robustly activated compared to the normal CD4^+^ and CD8^+^ tumor-infiltrating T-cells within the tumor immune microenvironment. Collectively, these data identify DNTs as an inflammatory DNT population in glioma characterized by inflammatory cytokine production, activation-associated transcriptional programs, and reduced features of exhaustion relative to conventional tumor-infiltrating T-cells.

### DNT-associated tumors exhibit an activated myeloid compartment enriched for inflammatory signaling

Given the inflammatory transcriptional profile observed in DNTs, we next investigated whether enrichment of this population altered the glioma myeloid compartment. Using immune-enriched scRNAseq from ΔCD4/Cd8^-/-^ and IgG/Cd8^+/+^ gliomas. TAMs from ΔCD4/CD8^-/-^ tumors demonstrated a striking transcriptional shift compared to TAMs from IgG/Cd8^+/+^ tumors (Figure 3a). Differential expression and pathway analyses revealed enrichment of chemokine signaling, CXCR-associated pathways, and inflammatory programs associated with myeloid activation in ΔCD4/Cd8^-/-^ TAMs (Figure 3b-c). In contrast, TAMs from IgG/Cd8^+/+^ tumors upregulated immunosuppressive pathways including PD-1/PD-L1 signaling and JAK-STAT signaling, both of which are associated with tumor tolerance and immune dysfunction (Figure 3b-c). Single-cell gene set enrichment analysis further demonstrated that ΔCD4/CD8^-/-^TAMs were enriched for inflammatory signaling, FasL-associated pathways, neutrophil recruitment, immune complex clearance, and cytotoxicity, whereas IgG/Cd8^+/+^ TAMs exhibited transcriptional programs associated with tolerance to tumor cells and impaired phagocytic function (Figure 3d).

**Figure 3:**
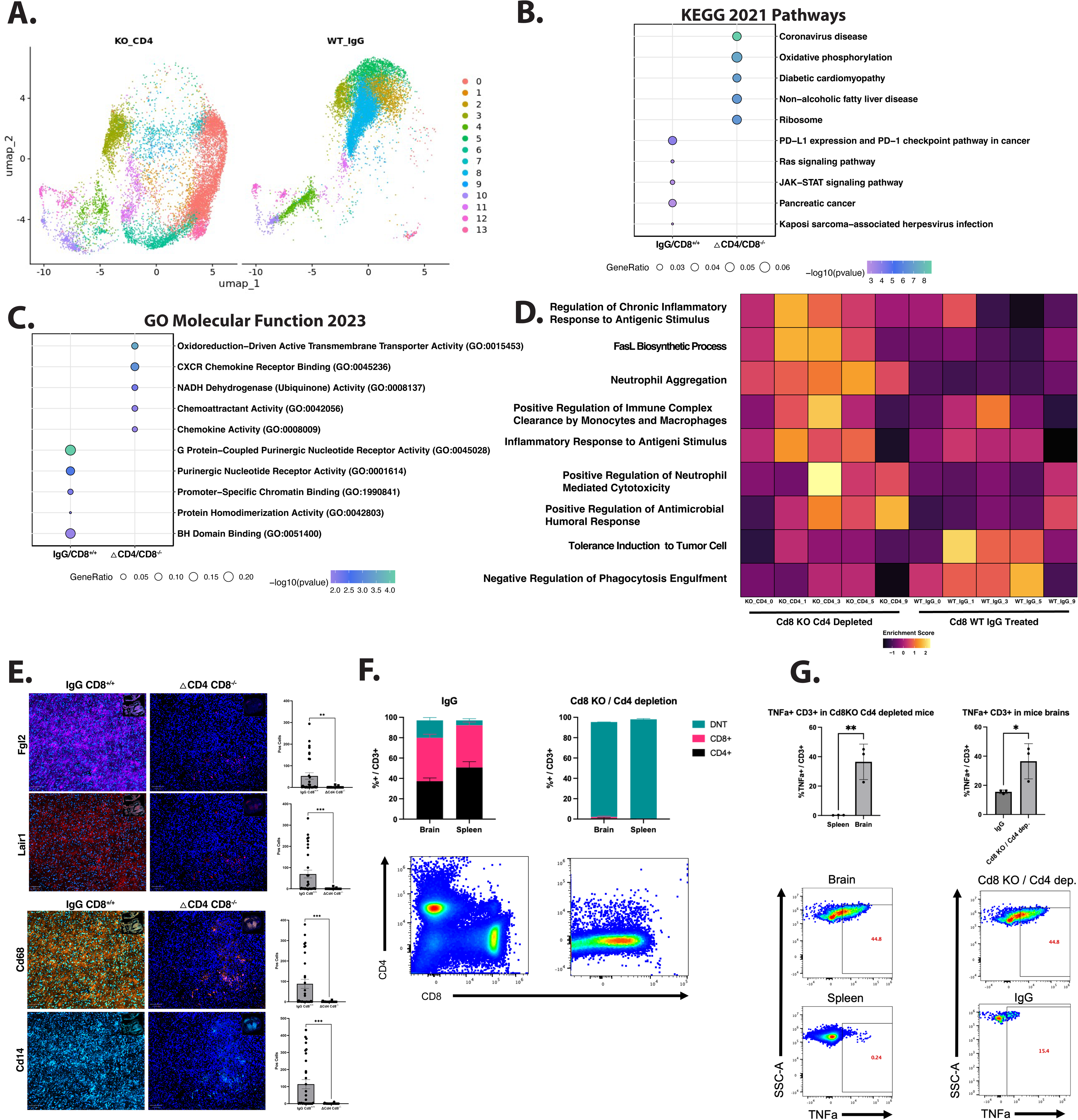
scRNAseq identifies activated myeloid population in ΔCD4/Cd8^-/-^. (A). Transcriptional differences between ΔCD4/Cd8^-/-^ and IgG/Cd8^+/+^ TAMs (B). Gene ontologies for DEGs from each group for Molecular Function and KEGG pathways. (D). single cell Gene Set Enrichment Analysis for different TAMs clusters by experiment. (E). Increased expression of immunosuppressive markers within tumor associated macrophages in IgG/Cd8^+/+^ compared ΔCD4/Cd8^-/-^ (Fgl2: p-val = 0.002, t-test; Lair1: p-val = 0.001, t-test). Multiplex fluorescence imaging for myeloid populations demonstrating increased tumor associated macrophages in IgG/Cd8^+/+^ compared ΔCD4/Cd8^-/-^ (Cd68: p-val = 0.005, t-test; Cd14: p-val = 0.002, t-test). (F). Flow cytometry data showing successful functional depletion of CD4 and CD8 positive T cells in brains and spleens of ΔCD4/Cd8^-/-^ tumor bearing mice compared to brains and spleens of tumor bearing IgG/Cd8^+/+^ mice. (G). Flow cytometry data showing increase in TNFα expression within CD3+ T cells from the brain of ΔCD4/Cd8^-/-^ tumor bearing mice compared to the spleen of ΔCD4/Cd8^-/-^ tumor bearing mice (*p <* 0.01) as well as increase in TNFα expression amongst CD3+ T cells from ΔCD4/Cd8^-/-^ tumor bearing mice when compared to CD3+ T cells from the brains of tumor bearing IgG/Cd8^+/+^ mice (*p <* 0.01)

Multiplex immunofluorescence demonstrated reduced infiltration of Cd68^+^ and Cd14^+^ TAMs in ΔCD4/Cd8^-/-^ tumors compared to IgG/Cd8^+/+^ tumors (Figure 3e). Moreover, TAMs within ΔCD4/CD8^-/-^tumors exhibited reduced expression of the immunosuppressive markers Fgl2 and Lair1, suggesting a less suppressive myeloid phenotype (Figure 3e). To further define communication networks between tumor-infiltrating lymphocytes (TILs) and TAMs, we performed CellChat analysis (Supp Figure 3). ΔCD4/CD8^-/-^tumors demonstrated enhanced TNFα-mediated signaling between TILs and TAMs, along with enrichment of innate immune activation pathways including GAS and CypA signaling. In contrast, IgG/CD8^+/+^ tumors preferentially upregulated suppressive signaling pathways including Cd86-Ctla4, MIF, Galectin, PTN, and TGFβ signaling. To validate protein level changes in TNFα, we evaluated TNFα expression by flow cytometry in tumor-bearing mice. Consistent with our single cell sequencing data, we found that ΔCD4/CD8^-/-^ contained predominantly DNTs in both the brain and the spleen (Figure 3f). DNTs isolated from ΔCd4/Cd8^-/-^ mice tumor-bearing brains demonstrated increased TNFα expression compared to splenic DNTs from ΔCd4/Cd8^-/-^ mice tumor-bearing mice (Figure 3g). Furthermore, DNT CD3^+^ T-cells isolated from ΔCD4/CD8^-/-^ tumors, exhibited greater TNFα expression than CD3^+^ T-cells from brain IgG/Cd8^+/+^ tumor bearing mice (Figure 3g). Together, these findings suggest that DNT-derived inflammatory signaling contributes to reprogramming the glioma myeloid compartment toward a more activated state.

### Human DNTs directly suppress glioma stem cells and promote inflammatory macrophage remodeling

To determine whether human DNTs possess functional antitumor activity against glioma, we isolated total T-cells (PanT) followed by DNTs from healthy donors and patients with newly diagnosed treatment naive GBMs using magnetic bead isolation followed by activation and expansion for 1 week in vitro. Purity of PanTs and DNTs was confirmed using flow cytometry (Supp. Fig 4). DNTs from healthy donors were predominately TCR γδ negative, CD25+ and variably expressed CD161 consistent with an activated non-conventional T-cell phenotype (Supp. Fig 4).

**Figure 4:**
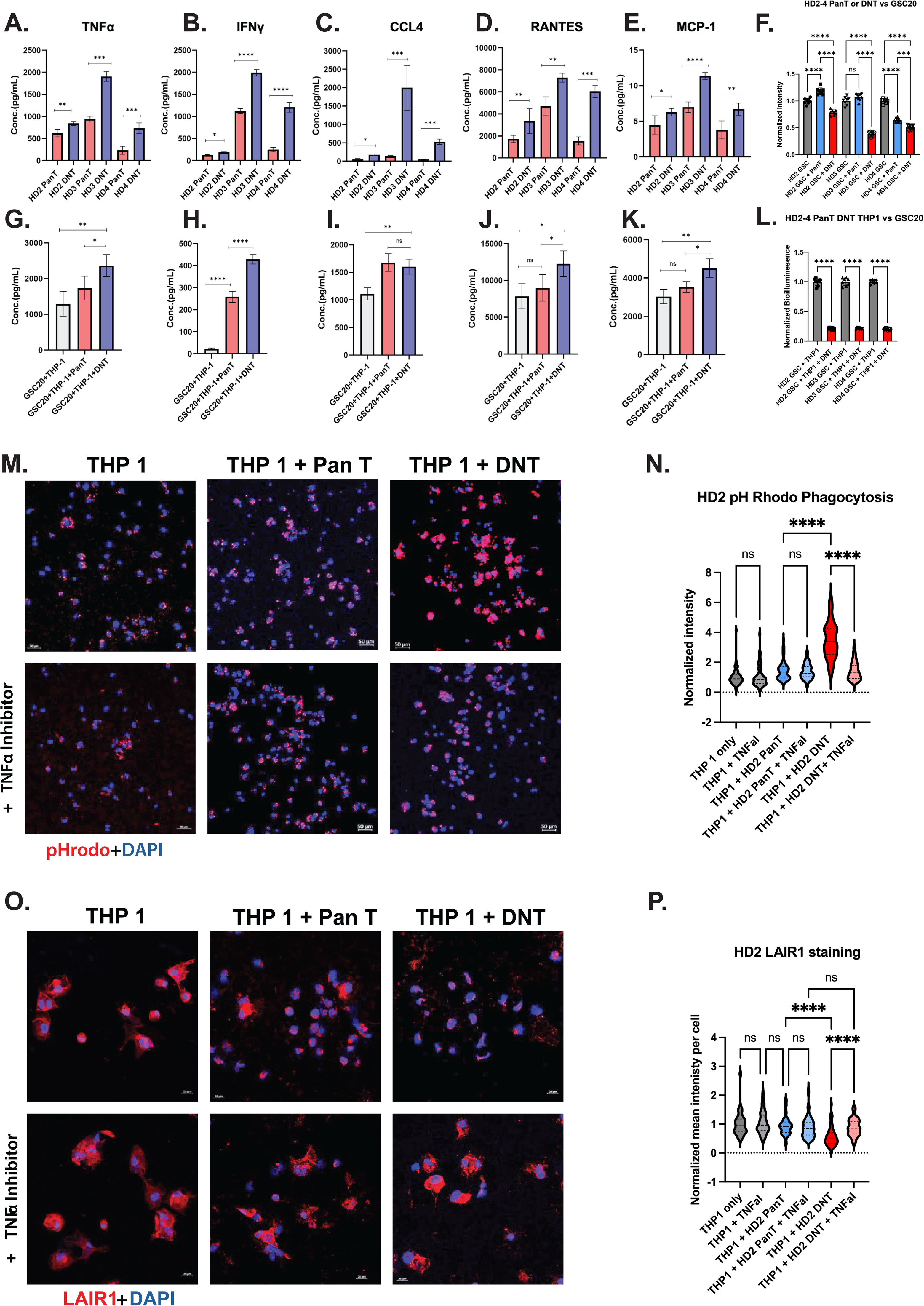
Human DNTs suppress glioma cell viability and promote macrophage phagocytic activity with reduced LAIR1 expression. (A-E). Cytokine profiling of healthy donor-derived DNTs and PanT cells cocultured with GSC20 glioma stem cells. Concentrations of TNFα (A), IFNγ (B), CCL4 (C), RANTES (D), and MCP-1 (E) were quantified by multiplex cytokine analysis. Data are shown for three independent healthy donors (HD2–HD4). Symbols represent technical replicate measurements; horizontal bars indicate mean ± s.d. (F). Luciferase-based assessment of GSC20 viability following coculture with PanT cells or DNTs from healthy donors (HD2–HD4). Tumor viability is shown as normalized relative bioluminescence (a.u.). Each symbol represents an independent experimental replicate. Horizontal bars indicate mean ± s.d. Cytokine profiling of GSC20 in the presence of healthy donor PanT or DNTs. Concentrations of TNFα (G), IFNγ (H), CCL4 (I), RANTES (J), and MCP-1 (K) were quantified by multiplex cytokine analysis. Data are shown as individual replicate measurements with mean ± s.d. (L). Luciferase-based assessment of GSC20 viability in the presence of THP1 macrophages. GSC20 cells were cocultured with THP1 macrophages alone or with DNTs DNTs from healthy donors (HD2–HD4). Tumor viability is shown as normalized relative bioluminescence (a.u.). Each symbol represents an independent experimental replicate. Horizontal bars indicate mean ± s.d. (M-N). Representative images and quantification of THP1 macrophage phagocytosis using pHrodo-labeled targets. THP1 cells were cultured alone, with PanT cells, or with DNTs in the presence or absence of TNFα inhibition. Following incubation with pHrodo substrate, fluorescence intensity was quantified by ImageJ and normalized to THP1-only controls. Each point represents an individual analyzed cell pooled from seven imaged fields. Horizontal bars indicate mean ± s.d. (O-P). Representative images and quantification of LAIR1 expression in THP1 macrophages cultured alone, with PanT cells, or with DNTs in the presence or absence of lenalidomide administration. Following coculture, non-adherent T cells were removed, adherent THP1 cells were stained for LAIR1 and DAPI, and fluorescence intensity was quantified by ImageJ. Data were normalized to THP1-only controls. Each point represents an individual analyzed cell pooled from seven imaged fields. Horizontal bars indicate mean ± s.d. For panels b–f, statistical significance was determined using two-tailed unpaired Welch’s t-tests for pairwise comparisons. For cytokine analyses, individual replicate measurements were compared between indicated groups. For phagocytosis and LAIR1 assays, comparisons were performed between predefined experimental conditions as indicated in the figure. Significance is denoted as \**P < 0.05,** P < 0.01,\*\*\** P < 0.001, ****P < 0.0001; ns, not significant.

To characterize the cytokine secretion patterns of PanT and DNT cells derived from healthy donor PBMCs, we performed a Luminex multiplex assay following co-culture with GSC20 glioma stem cells. Across all donors, DNT cells exhibited significantly higher secretion of pro-inflammatory cytokines compared to donor-paired PanT cells, including TNF-α, IFN-γ, and CCL4, as well as chemokines RANTES and MCP-1 (Figure 4a-e). These increases were consistent across donors, with multiple comparisons reaching high statistical significance (p < 0.01 to p < 0.0001). The data indicate that DNT cells possess a heightened inflammatory and chemotactic profile relative to PanT cells. We also noticed elevated levels of Th1-associated chemokine, CXCL9 (Mig) in 2/3 DNT samples compared to PanT cells (Supp Figure 5a). Because CXCL9 is a downstream responder gene strictly dependent on IFNγ/STAT1 activation, this differential upregulation demonstrates a donor-dependent CXCL9 secretion in THP1 co-culture compared to PanT. To further assess inflammatory cytokine production, we measured IL6 and IL8 levels under the same experimental conditions (Supp Figure 5b-c). In donor-derived PanT and DNT cultures, IL6 secretion was modest, with a slight increase observed in HD4 PanT compared to DNT (p < 0.05), while IL8 levels remained largely unchanged across donors (Supp Figure 5c).

**Figure 5:**
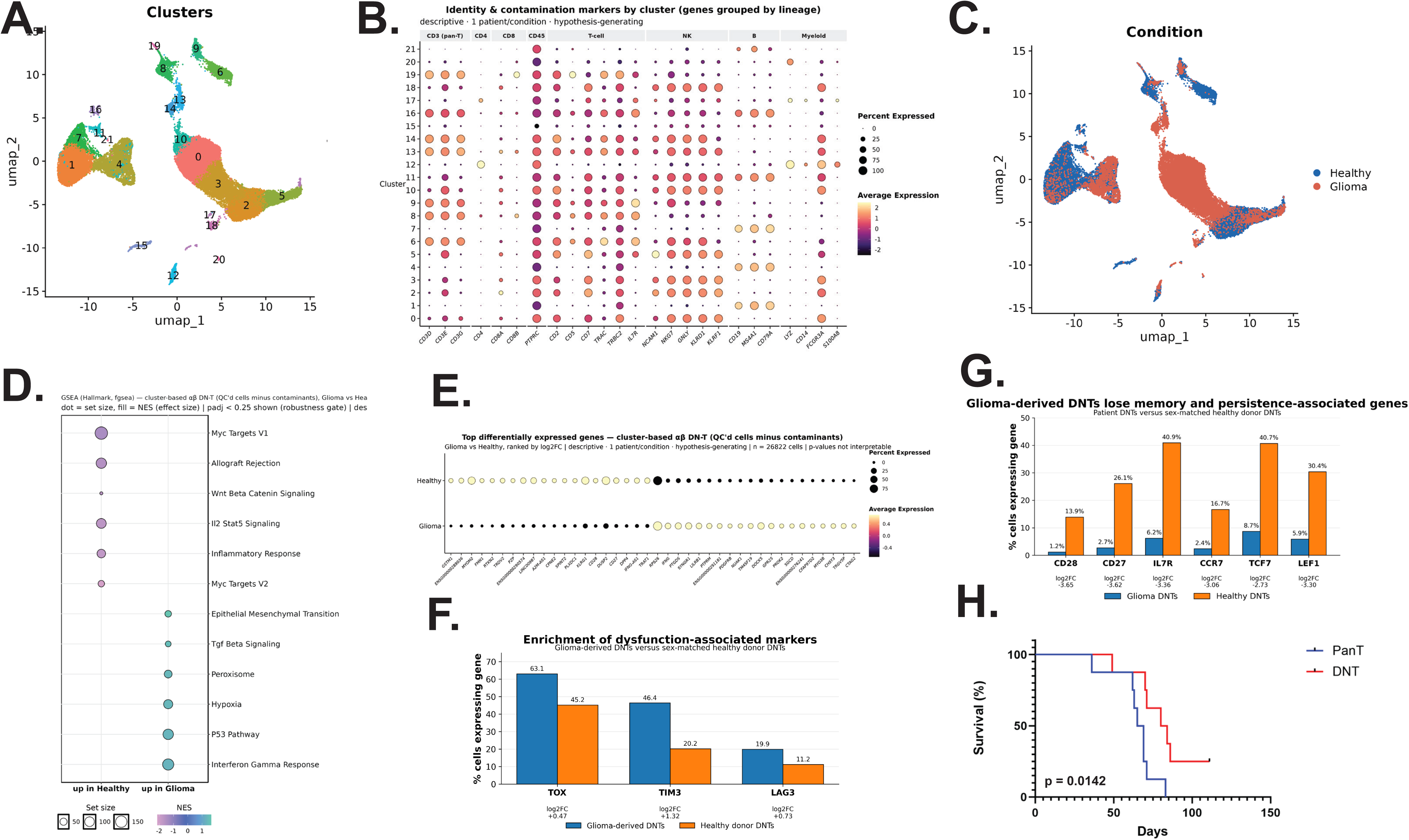
Glioma patient-derived DNTs exhibit a dysfunction-associated transcriptional phenotype, supporting evaluation of healthy donor-derived DNTs as an allogeneic cellular therapy. (A-C). Exploratory single-cell RNA-sequencing analysis of circulating DNTs isolated from one healthy donor and one patient with newly diagnosed glioblastoma prior to surgery, corticosteroid exposure, chemotherapy, or radiation. DNTs were isolated by fluorescence-activated cell sorting as CD45⁺CD3⁺CD4⁻CD8⁻ lymphocytes. (A). UMAP visualization demonstrating transcriptionally distinct DNT clusters identified by unsupervised clustering. (B). Dot plot showing expression of lymphoid lineage and identity markers across clusters, used to characterize DNT populations and assess potential cellular contamination. Dot size represents the percentage of cells expressing each gene and color represents average expression. (C). UMAP visualization of DNTs according to sample origin (healthy donor versus glioma patient), demonstrating differential distribution of cells across transcriptional states. (D). Hallmark gene-set enrichment analysis comparing healthy donor- and glioma patient-derived DNTs. Selected differentially enriched pathways are shown; normalized enrichment score (NES) indicates relative enrichment in healthy donor- or glioma patient-derived DNTs, with dot size reflecting gene-set size. (E). Differential gene-expression analysis comparing glioma patient- and healthy donor-derived DNTs following exclusion of clusters identified as potential non-DNT contaminants. Dot size represents the percentage of cells expressing each gene and color represents average expression. (F). Expression of dysfunction-associated genes in glioma patient-versus healthy donor-derived DNTs. Glioma patient-derived DNTs demonstrated increased frequencies of cells expressing TOX, HAVCR2 (TIM3), and LAG3. Bars represent the percentage of cells expressing each gene; corresponding log2 fold changes are displayed. (G). Expression of genes associated with T-cell memory, homeostatic maintenance, co-stimulation, and progenitor-like states. Compared with the healthy donor, glioma patient-derived DNTs demonstrated reduced frequencies of cells expressing CD28, CD27, IL7R, CCR7, TCF7, and LEF1. Bars represent the percentage of cells expressing each gene; corresponding log2 fold changes are displayed. Given the single individual analyzed per condition, panels A–G are descriptive and hypothesis-generating and were not used to infer population-level differences. (H). Kaplan–Meier survival analysis of mice bearing orthotopic GSC20 glioma xenografts treated with weekly intracranial administration of healthy donor-derived PanT cells or DNTs following tumor engraftment. Healthy donor-derived DNT treatment significantly prolonged survival compared with PanT treatment (log-rank [Mantel–Cox] test, P = 0.0142). **Abbreviations:** DNT, CD3⁺CD4⁻CD8⁻ double-negative T cell; PanT, pan-T cell; GBM, glioblastoma; GSC, glioma stem cell; UMAP, uniform manifold approximation and projection; NES, normalized enrichment score; TIM3, T-cell immunoglobulin and mucin-domain containing-3 (HAVCR2).

Next, we evaluated the direct antitumor activity of DNTs against glioma stem cells. Healthy donor-derived DNTs or PanT-cells were cocultured with luciferase-expressing GSC20 cells, and tumor viability was quantified by bioluminescence. DNTs consistently induced greater suppression of tumor viability than PanT-cells across all three independent healthy donor samples (Figure 4f). Similar findings were observed using a second glioma stem cell line, GSC6-27 (Supp Figure 6a), demonstrating that the antitumor activity of DNTs is not restricted to a single tumor model.

Because our murine studies suggested that DNTs may influence the myeloid compartment for tumor survival, we next examined whether DNTs could enhance macrophage-mediated tumor suppression. PMA-induced THP-1 macrophage-like cells were cocultured with GSC20 cells in the presence of either PanT-cells or DNTs. Consistent with the role for DNT-mediated myeloid modulation, the GSC20+THP-1 co-culture system demonstrated both PanT and DNT cells contributed to cytokine release, but DNT co-cultures demonstrated markedly higher levels of TNFα, IFNγ, RANTES, and MCP-1 compared to PanT co-cultures (Figure 4g-h,j-k). This suggests that DNT cells maintain robust effector functions even in the presence of tumor and macrophage-derived signals. Interestingly, CCL4 secretion showed donor-dependent variability, with some comparisons not reaching statistical significance, indicating selective cytokine modulation rather than uniform activation (Figure 4i). In the tumor co-culture system, CXCL9 and IL6 production were significantly elevated in the presence of PanT and DNT cells compared to GSC20+THP-1 alone (p < 0.01–0.001), suggesting that T cell engagement amplifies macrophage-driven IL6 release (Supp Figure 5d-e). Similarly, IL8 concentrations increased significantly upon co-culture with PanT and DNT cells (p < 0.05–0.01), indicating enhanced pro-inflammatory signaling and potential recruitment of neutrophils or other innate immune cells (Supp Figure 5f). DNT-containing cultures demonstrated significantly greater suppression of tumor viability than macrophage cultures containing PanT-cells across all three healthy donor samples when evaluating the same luciferase-based paradigm for GSC tumor viability (Figure 4l).

Whether cultured in isolation or co-cultured with THP1 cells, TNFα emerged as one of several inflammatory mediators that could contribute to macrophage effector function. THP1 macrophages were cocultured with PanT-cells or DNTs and phagocytic activity was quantified using pHrodo fluorescence imaging. DNT-exposed macrophages demonstrated significantly greater phagocytic activity than macrophages cultured alone or with PanT-cells (Figure 4m-n, Supp Figure 6b-c). Importantly, pharmacologic inhibition of TNFα production with lenalidomide attenuated this increase in phagocytosis, implicating TNFα as one contributor to DNT-mediated macrophage activation. Similar trends were observed across independent healthy donor experiments (Supp Figure 6d-e).

We next examined expression of the inhibitory myeloid receptor LAIR1, which was reduced in tumor-associated macrophages from murine ΔCd4/Cd8^−/−^ gliomas (Figure 3e). Coculture with DNTs resulted in significantly decreased LAIR1 expression in THP1 macrophages compared with PanT-containing cultures or macrophages alone (Figure 4o-p, Supp Figure 6f-g). Lenalidomide treatment restored LAIR1 expression toward baseline levels (Figure 4p, Supp Figure f-g). Together, these findings associate DNT exposure with increased macrophage phagocytic activity and reduced LAIR1 expression, both of which were sensitive to lenalidomide treatment. This effect was reproducible across independent healthy donor experiments (Supp Figure 6f-g).In contrast to healthy donor-derived DNTs, patient-derived T-cell populations demonstrated limited antitumor activity and failed to enhance macrophage phagocytosis (Supp Figure 6h-i). Although preliminary, these findings are consistent with evidence in the literature of systemic immune dysfunction in patients with glioblastoma and support the potential advantages of healthy donor-derived DNT products as an off-the-shelf cellular therapy platform.

Collectively, these data demonstrate that human DNTs suppress glioma growth through both direct and indirect mechanisms. In addition to directly impairing glioma stem cell viability, DNTs remodel macrophage inflammatory signaling, enhance phagocytic activity, and reduce expression of the inhibitory receptor LAIR1, supporting a broader role for DNTs in shaping the glioblastoma immune microenvironment.

### Glioma patient-derived DNTs exhibit features of dysfunction, supporting evaluation of healthy donor-derived DNTs as an allogeneic cellular therapy

To investigate whether DNTs isolated from patients with glioblastoma exhibit transcriptional features that may limit their therapeutic utility, we performed an exploratory single-cell RNA-sequencing analysis of DNTs isolated from a single sex matched female healthy donor and a single female patient with newly diagnosed glioblastoma. The patient was treatment-naive at the time of peripheral blood collection and had not received corticosteroids, surgery, chemotherapy, or radiation. DNTs were isolated by fluorescence-activated cell sorting as CD45^+^CD3^+^CD4^-^CD8^-^lymphocytes and subjected to single-cell transcriptomic analysis for just these two individuals. Because this analysis represents one individual per condition, these data were considered hypothesis-generating rather than representative of population-level differences.

Unsupervised clustering identified transcriptionally heterogeneous DNT populations in both samples (Figure 5a-c). Pathway-level analysis demonstrated differential enrichment of transcriptional programs between healthy donor- and glioma patient-derived DNTs (Figure 5d), while differential gene-expression analysis further suggested substantial differences in DNT cell state (Figure 5e). In particular, glioma patient-derived DNTs demonstrated increased expression of dysfunction-associated genes, including TOX, HAVCR2 (TIM3), and LAG3, compared with healthy donor-derived DNTs (Figure 5f). Conversely, genes associated with T-cell memory, homeostatic maintenance, and progenitor-like states - including CD28, CD27, IL7R, CCR7, TCF7, and LEF1 - were markedly reduced in patient-derived DNTs (Figure 5g). Together, these exploratory findings suggest that circulating DNTs from this treatment-naive glioblastoma patient occupied a more differentiated and dysfunction-associated transcriptional state relative to the healthy donor comparator.

These transcriptional findings were consistent with our observations in which patient-derived DNTs were difficult to expand *ex-vivo* and, when expansion was successful, demonstrated limited antitumor activity and failed to enhance macrophage phagocytosis compared with healthy donor-derived DNTs (Supp Figure 6). Given these observations, together with prior evidence supporting the feasibility of allogeneic DNT-based cellular therapy, we next asked whether healthy donor-derived DNTs could provide therapeutic activity against glioblastoma in vivo. SCID mice underwent cranial bolt placement followed by intracranial implantation of 2.5 × 10⁵ luciferase-expressing GSC20 glioma stem cells. Tumor engraftment was confirmed one week later by bioluminescence imaging, after which mice received weekly intracranial administration of either PanT-cells or DNTs through the implanted cranial bolt. DNT-treated mice demonstrated significantly prolonged survival compared with mice treated with PanT-cells (Figure 5h). Median overall survival increased from 67 days in the PanT-treated group to 82 days in the DNT-treated group, representing a 15-day survival benefit. Survival analysis demonstrated a significant advantage for DNT therapy by log-rank testing (p = 0.0142). These findings demonstrate that unmodified healthy donor-derived DNTs provide a survival advantage over PanT cells in an intracranial patient-derived glioblastoma model, without CAR engineering or introduction of an artificial target antigen into the tumor. Whether myeloid modulation contributes to this therapeutic effect remains to be determined.

## Discussion

Our study began with the unexpected observation that CD4 depletion in CD8-deficient mice prolonged survival following glioma induction. Investigation of the remaining tumor-infiltrating lymphocytes identified an inflammatory DNT population accompanied by changes in tumor-associated myeloid cells. Complementary human studies demonstrated that unmodified healthy donor-derived DNTs suppress glioma stem cell viability, alter macrophage phenotype and phagocytic activity, and prolong survival in an orthotopic xenograft model. These findings identify both tumor-directed activity and myeloid-modulating properties of DNTs, while leaving open the extent to which each contributes to tumor control in vivo.

Unexpectedly, simultaneous elimination of conventional CD4⁺ and CD8⁺ T-cells resulted in a significant survival advantage in murine glioma and revealed a previously unappreciated population of DNTs. Single-cell transcriptomic analysis demonstrated that these DNTs possessed a transcriptional profile distinct from conventional tumor-infiltrating T-cells, characterized by activation-associated and inflammatory programs rather than the exhausted phenotype observed in conventional lymphocytes. DNTs demonstrated increased expression of inflammatory mediators, including TNF-associated pathways, together with activation markers such as CD69, CD226, and IL7R, while conventional T-cells preferentially expressed canonical exhaustion-associated genes. These findings suggest that DNTs represent a functionally distinct immune population within gliomas rather than simply residual conventional T-cells lacking CD4 or CD8 expression.

Although DNTs have previously been described in autoimmune disease, transplantation biology, and hematologic malignancies, their role in glioblastoma remains poorly understood^16–25^. Prior studies have demonstrated that DNTs can exert direct anti-tumor activity through mechanisms including Fas-FasL interactions, perforin-mediated killing, and inflammatory cytokine production^23–25,27–30^. Consistent with these observations, healthy donor-derived DNTs demonstrated robust anti-tumor activity against glioma stem cells in vitro and significantly prolonged survival in an orthotopic xenograft model. Together, these findings support the emerging concept that DNTs represent a therapeutically relevant effector population capable of mediating anti-tumor immunity in otherwise immunologically resistant tumors.

Prior studies established the feasibility of healthy donor-derived allogeneic CAR-DNT therapy, including CD19-directed activity against B-cell acute lymphoblastic leukemia and lung tumors engineered to express CD19^31^. A recent conference abstract also reported B7-H3-directed CAR-DNT activity in glioblastoma models and CAR-independent cytotoxicity against tumor core-derived cells^32^. Our findings extend this work by investigating the inflammatory phenotype of DNTs persisting in gliomas after conventional T-cell removal, their association with altered myeloid states, and the effects of unmodified human DNTs on macrophage function. The therapeutic activity observed with unmodified donor-derived DNTs further supports evaluating their native functions alongside approaches that introduce antigen-specific CAR recognition.

A central finding of this study is that DNTs appear to influence glioma-associated myeloid populations. Both single-cell transcriptomic analysis and multiplex immunofluorescence demonstrated substantial differences within the myeloid compartment of DNT-enriched tumors, including reduced expression of immunosuppressive markers and enrichment of inflammatory signaling pathways. These observations were further supported by in vitro coculture studies in which DNTs altered macrophage-associated cytokine production and enhanced macrophage-mediated antitumor activity. Mechanistically, we identified a broad inflammatory cytokine program associated with DNTs that included TNFα, IFNγ, and CCL4. Among these mediators, TNFα emerged as one experimentally tractable pathway linking DNTs to macrophage function. Lenalidomide attenuated DNT-associated macrophage phagocytosis and restored expression of the inhibitory receptor LAIR1, suggesting that TNFα contributes to specific aspects of macrophage activation. LAIR1 is an inhibitory collagen-binding receptor that has been implicated in immune suppression across multiple tumor types and has recently been established as a potential regulator of myeloid cell function within glioblastoma^33–36^. The reduction in myeloid LAIR1 expression for DNTs in our studies raises the possibility that DNTs promote a less suppressive macrophage phenotype through modulation of inhibitory receptor signaling. DNTs produced multiple inflammatory mediators, and the observed biological effects are unlikely to be attributable to a single cytokine alone. We therefore view TNFα and LAIR1 as one component of a broader inflammatory network through which DNTs influence myeloid cell behavior. Future studies will be required to define the relative contributions of additional pathways, including IFNγ and chemokine-associated signaling networks, within the glioblastoma microenvironment. Collectively, these findings suggest that DNTs may function not only as direct anti-tumor effectors but also as regulators of the glioblastoma immune microenvironment.

The findings presented here may also have implications for the development of next-generation cellular therapies for glioblastoma. Most CAR-T-cell strategies currently rely on conventional T-cells, which are frequently characterized by exhaustion and functional impairment in patients with glioblastoma^37–39^. In contrast, DNTs exhibited a less exhausted phenotype, retained robust inflammatory activity, and demonstrated the capacity to influence both tumor cells and macrophages. These characteristics raise the possibility that DNTs may provide a favorable cellular platform for engineering adoptive immunotherapies. Importantly, prior studies have demonstrated the feasibility of generating CAR-engineered DNTs with potent antitumor activity in hematologic malignancies, supporting the translational potential of this approach^30,31^. Although CAR-modified DNTs were not evaluated in the present study, our findings suggest that combining antigen-specific targeting with the intrinsic inflammatory and myeloid-remodeling properties of DNTs may represent a promising strategy for future glioblastoma immunotherapy development. An additional finding of this study was the limited expansion and functional activity observed in patient-derived DNT cultures. Expansion was successful in only a single patient sample, and these cells demonstrated substantially reduced activity compared with healthy donor-derived DNTs. Although exploratory, these findings are consistent with extensive evidence demonstrating systemic immune dysfunction and T-cell exhaustion in patients with glioblastoma^40^. From a translational perspective, these observations may support the development of healthy donor-derived DNT products as an off-the-shelf cellular therapy platform^22,25,31^. Such an approach is particularly attractive given prior reports demonstrating minimal graft-versus-host toxicity associated with DNT-based therapies in hematologic malignancies^25,30,31^.

## Limitations

Several limitations should be acknowledged. First, mechanistic studies were performed using THP1-derived macrophages rather than primary human macrophages. While THP1 cells provide a reproducible platform for studying macrophage biology, future studies incorporating primary monocyte-derived macrophages and patient-derived myeloid populations will be important for validation. Second, the patient-derived DNT analyses were limited by the inability to consistently expand DNTs from glioblastoma patient samples. Third, although TNFα inhibition attenuated macrophage phagocytosis and restored LAIR1 expression, these experiments do not establish TNFα as the sole mediator of DNT function. Furthermore, lenalidomide has other immunomodulatory effects beyond suppression of TNFα secretion, potentially confounding some of the results. Finally, additional studies will be required to determine whether DNT-mediated myeloid remodeling directly accounts for the survival benefit observed in vivo.

## Conclusion

In conclusion, we identify DNTs as a distinct inflammatory lymphocyte population associated with improved survival and enhanced antitumor immunity in glioblastoma. Beyond their direct cytotoxic activity, DNTs appear capable of reshaping the glioma myeloid compartment through inflammatory signaling pathways that promote macrophage activation and reduce expression of inhibitory immune regulators. These findings establish DNTs as a previously underappreciated component of the glioblastoma immune response and support further development of DNT-based cellular immunotherapies for patients with glioblastoma.

## METHODS

### Cell culture

All cells were kept at 37 C with 5% CO2 in a humidified incubator. Chicken DF1 and RAW 264.7 Macrophages cells were cultured in DMEM (Corning Inc), supplemented with 10% FBS and 1% penicillin/streptomycin. Cells were maintained by trypsin passage when confluent. Patient derived glioma cells were cultured in DMEM/F12 50/50, 1X B27 Supplement (Invitrogen #17504-044), 20ng/mL EGF (Sigma #E9644), 20ng/mL bFGF (Sigma #F0291), 1% Pen/Strep solution. T-cells were cultured in RPMI 1640 with 1% Pen/Strep. These cells were maintained by trypsin passage every 2–3 days. THP-1 cell line was cultured in RPMI 1640 medium supplemented with 10% heat-inactivated fetal bovine serum and 1% penicillin/streptomycin. THP-1 cells were induced with 10-ng/mL phorbol-12-myristate-13-acetate (PMA, Sigma Chemical, St. Louis, MO, USA) for 24 h and later changed to PMA-free media for 48hrs-72 hrs. and used for further experiments.

### Analyzing TCGA expression data and survival analysis

We obtained the Cancer Genome Atlas high-grade glioma (TCGA-GBM) and TCGA-low-grade glioma (LGG) raw read counts and clinical data using the TCGAbiolinks R package (RRID:SCR_017683). To assess the gene set enrichment in human data, we performed ssGSEA using the GSVA R package (RRID:SCR_021058) using expression of CD4, CD8A, and CD8B. We then stratified the bulk RNA-seq data by the ssGSEA gene score into upper and lower quartiles to assign samples and compared survival using a Kaplan–Meier survival model and log-rank test using survival R package (RRID:SCR_021137).

### Animal Models

Tumor bearing mice were generated as previously described^15,26,41^. Briefly, postnatal day 1-2 transgenic mice with the TVA receptor under a nestin promoter (nTVA) were injected bilaterally in the frontal lobes with 1x10^4^ DF1 cells infected with RCAS-PDGFB and 2.5x10^4^ DF1 cells infected with RCAS-STAT3 to drive overexpression as indicated. Mice were either nTVA-Cd8^-/-^ or nTVA-Cd8^+/+^ for Cd8 T-cell knockout. For the survival study, mice were humanely euthanized according to Baylor College of Medicine IACUC guidelines after displaying symptoms of tumor burden such as hydrocephalus, hunched posture, ataxia, limb paralysis, or lethargy or at 90 days. The survival date was recorded for Kaplan-Meier Analysis. Both male and female mice were utilized for all experiments. The RCAS-PDGFB and RCAS-STAT3 model recapitulates histopathological features of high-grade glioma including macrophage infiltration^15,26,41^. For the Xenograft studies, GSC20 cells were implanted intracranially in 4–5-week-old NOD/SCID mice. 2.5 × 10^5^ cells were injected intracranially in each mouse and after one-week freshly cultured PanT/DNT cells (2.5x 10^5^) from PBMCs of a healthy donor were injected. A minimum of eight mice (4 male, 4 female) were used in each group. Mice that presented neurological symptoms (i.e. hydrocephalus, seizures, inactivity, and/or ataxia) or moribund were sacrificed, and brains were fixed in formalin and stained with H&E to confirm the presence of tumor and subject to IHC. All animal procedures were reviewed and approved by the Institutional Animal Care and Use Committee at the Baylor College of Medicine.

### Antibody Depletion

To generate mice deficient in CD4+ T-cells, RCAS-Ntva mice were treated with either anti-CD4 antibody (1.33 ug/uL, 200 ug total, InVivoMAb BE0003-1) or isotype control IgG at the same concentration and dose on postnatal day 10, 15, 20, 25, and 30. Depletion was confirmed via flow cytometry at P60.

### PBMC Extraction

Peripheral blood was collected in sodium heparin vacutainer collection tubes (BD #366480) from patients with glioma with their preoperative informed consent to participate in research under Institutional Review Board H35355 at Baylor College of Medicine. A density gradient was established using centrifugation and Histopaque®-1077 (Sigma-Aldrich #10771-500ML). The PBMC layer was selected, rinsed with sterile PBS, and collected for designated experiment.

### T-cell Isolation

Mice were humanely euthanized at designated timepoint and had their spleens dissected and digested using a collagenase mix. A density gradient was established using Percoll, and immune cells were selected. T-cells were isolated using a system of sequential magnetic bead selection (Pan-Tcell Isolation Miltenyi Biotec, Pan T-Cell Isolation Kit II (mouse: 130-095-130, human: 130-096-535), CD4 MicroBeads mouse, Miltenyi Biotec, (mouse: 130-117-043, human: 130-097-048)) and run through a magnetic column on a MACS magnetic separator.

### Culturing Pan-T and DNT cells

CD4 and CD8 cells depleted DNT and Pan-T cell enriched population isolated from healthy donor PBMCs were cultured on anti-CD3 and CD28 coated plates for 2 days in RPMI-1640 supplemented with 10% FBS. Cells were maintained in fresh RPMI-1640 media containing IL-7 and IL-15 from day 3 onwards. Purity of DNT cells were analyzed by flow cytometry using anti-human antibodies CD3, CD4 and CD8 and CD45.

### Flow cytometry analysis

Mouse spleens were harvested under sterile conditions and placed in cold PBS. Single-cell suspensions were prepared by mechanically dissociating the spleens through a 70 μm cell strainer, followed by red blood cell lysis using an ammonium chloride-based buffer. Cells were washed, counted, and plated into 96-well U-bottom plates (100 µL/well). Viability was assessed using LIVE/DEAD NIR dye (Invitrogen) for 30 min at room temperature in the dark. Cells were then stained with fluorochrome-conjugated antibodies against CD3, CD4, CD8, and CD45 (BioLegend, BD Biosciences). Antibodies were vortexed before use, diluted according to manufacturer recommendations, and added at 100 µL per well; volumes were adjusted based on the number of experimental and control wells, including single-stain and unstained controls for spectral compensation. After staining for 30 min at 4°C in the dark, cells were washed and resuspended in 200 µL FACS buffer, then maintained at 4°C in the dark until acquisition. Samples were acquired on a Cytek Aurora spectral flow cytometer, and data were analyzed using FlowJo software (BD Biosciences), with gating and compensation established using single-stain and fluorescence-minus-one (FMO) controls.

Peripheral blood mononuclear cells (PBMCs) were isolated and processed for flow cytometric analysis using fluorochrome-conjugated anti-human antibodies. Cells were washed and plated into 96-well U-bottom plates. Viability was assessed using LIVE/DEAD NIR dye (Invitrogen) for 30 min at room temperature in the dark. Surface staining was performed with fluorochrome-conjugated antibodies against CD3, CD4, CD8, CD25, CD45, TCR, CD161, and TNF (Supp Table 1, BioLegend, BD Biosciences, and Ah) for 30 min at 4°C in the dark. Single-stain and unstained controls were included for spectral compensation, and bead-based controls were prepared to standardize instrument settings. After staining, cells were washed and resuspended in 200 µL FACS buffer, then kept at 4°C in the dark until acquisition. Data was acquired on a Cytek Aurora spectral flow cytometer and analyzed using FlowJo software (BD Biosciences), with gating and compensation established using single-stain and fluorescence-minus-one (FMO) controls.

### Histology and Immunofluorescence

Tumor brains were collected for paraffin sectioning. Mouse brains were fixed in 4% paraformaldehyde overnight after perfusion. Brains were then dehydrated in 70% ethanol and embedded in paraffin by the Breast Center Pathology Core at Baylor College of Medicine. Samples were sectioned to a thickness of 10 um and deparaffinized with xylene, then rehydrated with successive decreasing concentration of ethanol. Hematoxylin and eosin were used for nuclear and cytoplasmic labeling. For immunofluorescence, antigen exposure was completed by incubating samples in 10 mM sodium citrate when necessary. Slides were blocked for 1 hour in serum and then incubated overnight with the primary antibody at 4C. The following day, slides were washed with PBS and stained with the corresponding conjugated secondary antibody. Slides were then counterstained with Hoecht, mounted, and sealed with antifade mounting media.

### Multiplexed Sequential Immunofluorescence (GEMM Model)

Sequential IF was performed using the COMET (Lunaphore) machine. 4 µm thick unstained Formalin Fixed and Paraffin Embedded (FFPE) tissue sections were mounted on charged slides prior to staining. Adjacent tissue sections received H&E staining to confirm tumor location. Primary antibodies used in this experiment at listed in Supplementary Table 2. All primary antibodies were paired with a corresponding secondary antibody based on original source: Alexa Fluor 555 (ThermoFisher, A32727, Dilution 1/200) and Alexa Fluor 647 (ThermoFisher, A32733, Dilution 1/400). Lastly, a DAPI counterstain was used for nuclear staining (ThermoFisher, 62248, Dilution 1/1500). Antigen retrieval from FFPE slides was completed using the PT Module (Epredia) with Dewax and HIER Buffer H (AR02, Lunaphore) for 90 minutes at 102°C, after which slides were submerged in Multistaining Buffer (BU06, Lunaphore) Staining cycles were 4min long for primary antibodies, with a 2min period for secondary antibodies. After each staining cycle, imaging was performed using Imaging Buffer (BU09, Lunaphore) to capture TRITC and Cy5 dye signal. A subsequent 2min elution step was performed with Elution Buffer (BU07-L, Lunaphore) with an additional 30sec quenching step utilizing Quenching Buffer (BU08-L, Lunaphore). Slide imaging produced a multi-stack OME-TIFF file as an alignment of individual cycle imaging. The OME-TIFF files contain a DAPI image and autofluorescence signal via TRITC and Cy5 channels, with a single fluorescent layer per marker. Multiplex imaging was conducted using a Lunaphore comet.

### Phagocytosis and LAIR1 analysis

THP1 cells and T-cells were incubated together for 48hrs followed by incubation with pH-Rodo E.coli (phagocytosis assay, Invitrogen, #P35361) for 1 hr then washed 3X with sterile PBS. For studies with lenalidomide (5 μM) was added to the media and incubated for 48hrs. Cells were then fixed with PFA and methanol and then counterstained with Hoecht and imaged using confocal microscopy. Intensity of Rodo or LAIR1 was quantified using ImageJ and normalized to all samples.

### Luciferase Assay

GSCs were plated in equal number in 64 well plates. ONE-Glo™ EX Luciferase Assay Buffer (Promega, E62A)was combined with ONE-Glo™ EX Luciferase Assay Substrate (Promega E33A) and cells were incubated for 10 minutes. Luciferase signaling was measured using a plate reader.

### Luminex analysis

The culture supernatant samples were collected from the PanT/DNT alone cultures and from GSC+THP-1 co-cultures with Pan-T/DNT cells, were aliquoted into 1.5ml centrifuge tubes, and stored at -80°C. Cytokine concentrations in the samples were measured in duplicate using Luminex bead-based assays (LAG3, Burlington, MA, USA) following the manufacturer’s instructions specific to each kit. Briefly, culture supernatant samples were thawed on ice, centrifuged at 14000 × g at 4 °C for 5 minutes, and then transferred to a 96-well plate. The samples were diluted 1:2 with assay buffer provided in the kit. In each well, a 25 µl of each sample was mixed with beads and incubated at 4°C on a shaker for 18 hours. Following incubation with the Biotin-Antibody Cocktail and subsequent Streptavidin-PE, samples were analyzed using the Bio-Plex 200 system (Bio-Rad, Hercules, CA, USA) and the Bio-Plex manager software 7.2 (Bio-Rad). Fifty beads per analyte were collected with a 60-second timeout set for plate analysis. Cytokine concentrations were calculated using a standard curve with a 5-parameter logistic-fitting method in Bio-Plex manager software, and the average values of the replicates were used for the final analysis.

### Single-cell RNA sequencing

Mice were humanely euthanized at designated timepoint and had tumor bearing regions of brain dissected and digested using a collagenase mix. A density gradient was established using Percoll, and immune cells were selected. Samples were run through a MACS dead cell removal kit (Miltenyi Biotec, 130-090-101) and red blood cell lysis kit (Miltenyi Biotec 130-094-183) for single cell isolation. Patient derived DNTs for sequencing were harvested from blood samples taken intraoperatively or from healthy donors. Peripheral blood mononuclear cells (PBMCs) from a healthy donor (JRT; HD2.5) or an adult GBM patient (PT2) were stained with 7-AAD, anti-human CD45-APC-Cy7, anti-human CD3-FITC, anti-human CD4-Spark Violet 538, and anti-human CD8-APC in 1X HBSS containing 2% FBS on ice for 20 minutes. DN T cells (7-AAD-CD45+CD3+CD4-CD8-singlets) were then sorted using a BD FACSDiscover S8 Sorter into 0.5 ml of 1X PBS containing 20% FBS in tubes coated with 100% FBS overnight in 4C. Sorted DN T cells were kept on ice and submitted to the core for single cell RNA-seq. Single cells were processed with the 10X Chromium 3′ Single-Cell Platform using the Chromium Single-Cell 3′ Library, Gel Bead, and Chip Kits (10X Genomics) following the manufacturer’s protocol. Briefly, approximately 5,000–15,000 cells were added to each channel of a chip to be partitioned into Gel Beads in Emulsion (GEMs) in the Chromium instrument, followed by cell lysis and barcoded reverse transcription of RNA in droplets. GEMs were broken, and cDNA from each single cell was pooled. Clean-up was performed using Dynabeads MyOne Silane Beads (ThermoFisher, 37002D). Subsequently, the cDNA was amplified and fragmented to optimal size before end repair, A-tailing, and adaptor ligation. Raw sequencing reads were processed using Cell Ranger v8.0.0 and mouse (GRCm39) reference genome. Cells were analyzed independently using Seurat^42^. Genes detected in fewer than three cells and cells expressing fewer than 300 or ≥5,000 or containing ≥10% mitochondrial transcripts, were excluded. Data were normalized, and the top 2,000 variable genes were selected for principal component analysis (30 PCs). Harmony was used to correct for sample-associated variation, followed by UMAP visualization and clustering at a resolution of 0.2 using 30 Harmony dimensions^43^. Cluster markers were identified using FindAllMarkers (adjusted P < 0.05, log2FC > 0.5, minimum detection fraction 0.2). Cell types were annotated using SCRAM reference-based predictions from neural, immune, developmental, and glioma datasets, with a prediction probability threshold of 0.9^44^.

### Statistical Analyses

Statistical details are listed for each experiment in the figure legends. All analyses were blinded to experimental conditions. Mice were randomly distributed to each group. Analysis was conducted using ImageJ/Fiji and visualized using GraphPad Prism and RStudio. For all immunofluorescence staining, each replicate represents an independent mouse with slides exposed to the same staining and confocal imaging conditions. Kaplan-Meier analysis was used for survival with log-rank and Gehan-Breslow-Wilcoxon values provided for significance. Data was assessed for normality using the Shapiro-Wilk test. Parametric tests were utilized for all normally distributed datasets unless indicated otherwise. Unpaired Student’s t-test or Welch’s unpaired T-test was used for comparing two groups. One-way analysis of variance (ANOVA) or Two-way ANOVA was used for groups of 3 or more with Tukey’s correction for multiple comparisons. Mann-Whitney U tests were used for non-normally distributed data.

## CONTRIBUTIONS

M.F.M., R.P., and G.R. are responsible for conception of this project, study design and interpretation of results. Y.Y. generated and maintained all RCAS tumor bearing mice with assistance from M.F.M. M.F.M and R.P. performed all in vitro experiments, immunofluorescence, microscopy experiments, and scRNAseq analysis with assistance from A.S.H, E.A.G, and Y.J.K. Multiplex immunofluorescence was performed by R.D, H.T.S.C, H.N, and analyzed by V.S. All flow cytometry experiments were completed by A.P. Project oversight and financial support was provided by B.D., A.B.H., and G.R.

