## Supplementary figures and images for "CD3⁺CD4⁻CD8⁻ (Double-Negative) T-cells Provide Inflammatory Signals That Influence Myeloid Cells and Enhance Survival in Glioblastoma"

### Supplemental Figure 1

**A.**

**IgG Cd8 $^{+/+}$**

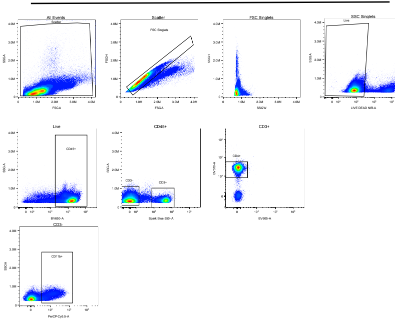


**B.**

**△Cd4 Cd8 $^{-/-}$**

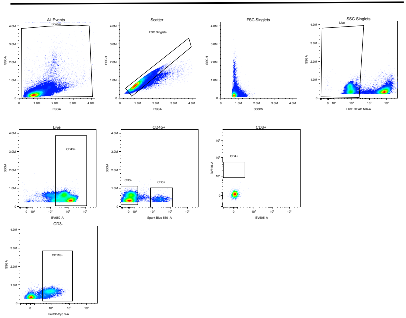

### Supplemental Figure 4

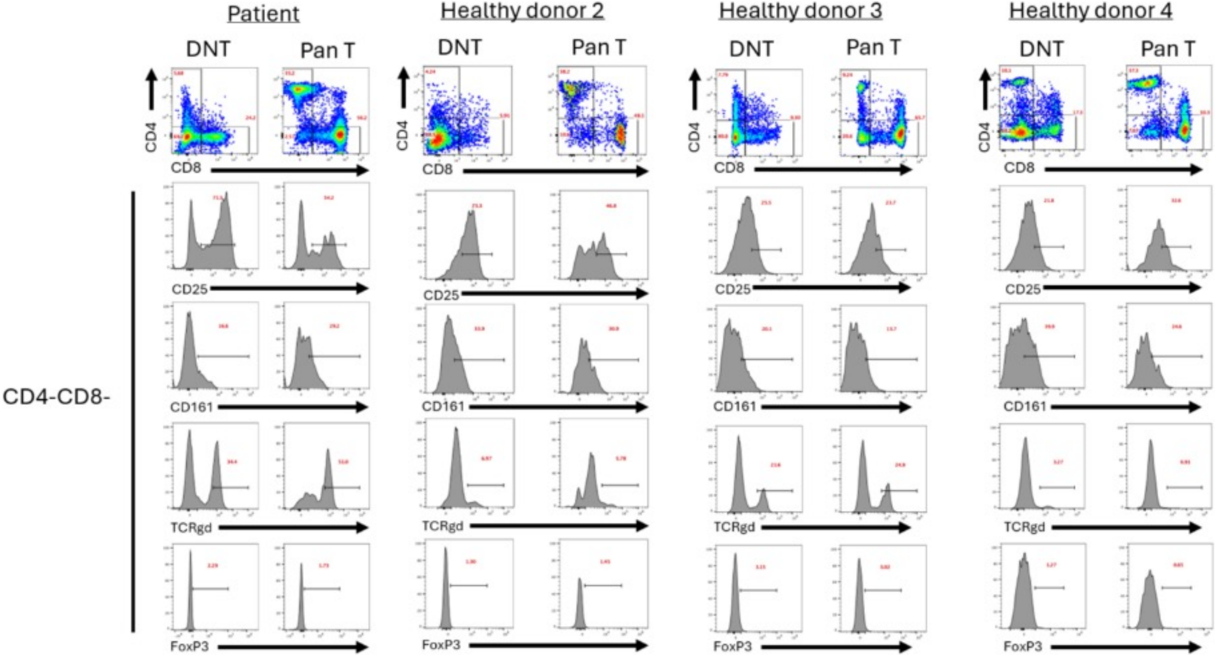

Patient
Healthy donor 2
Healthy donor 3
Healthy donor 4
DNT
Pan T
CD4
CD8
CD25
CD161
TCRgd
FoxP3
CD4-CD8-
