## Supplemental Figure 2 for "CD3⁺CD4⁻CD8⁻ (Double-Negative) T-cells Provide Inflammatory Signals That Influence Myeloid Cells and Enhance Survival in Glioblastoma"

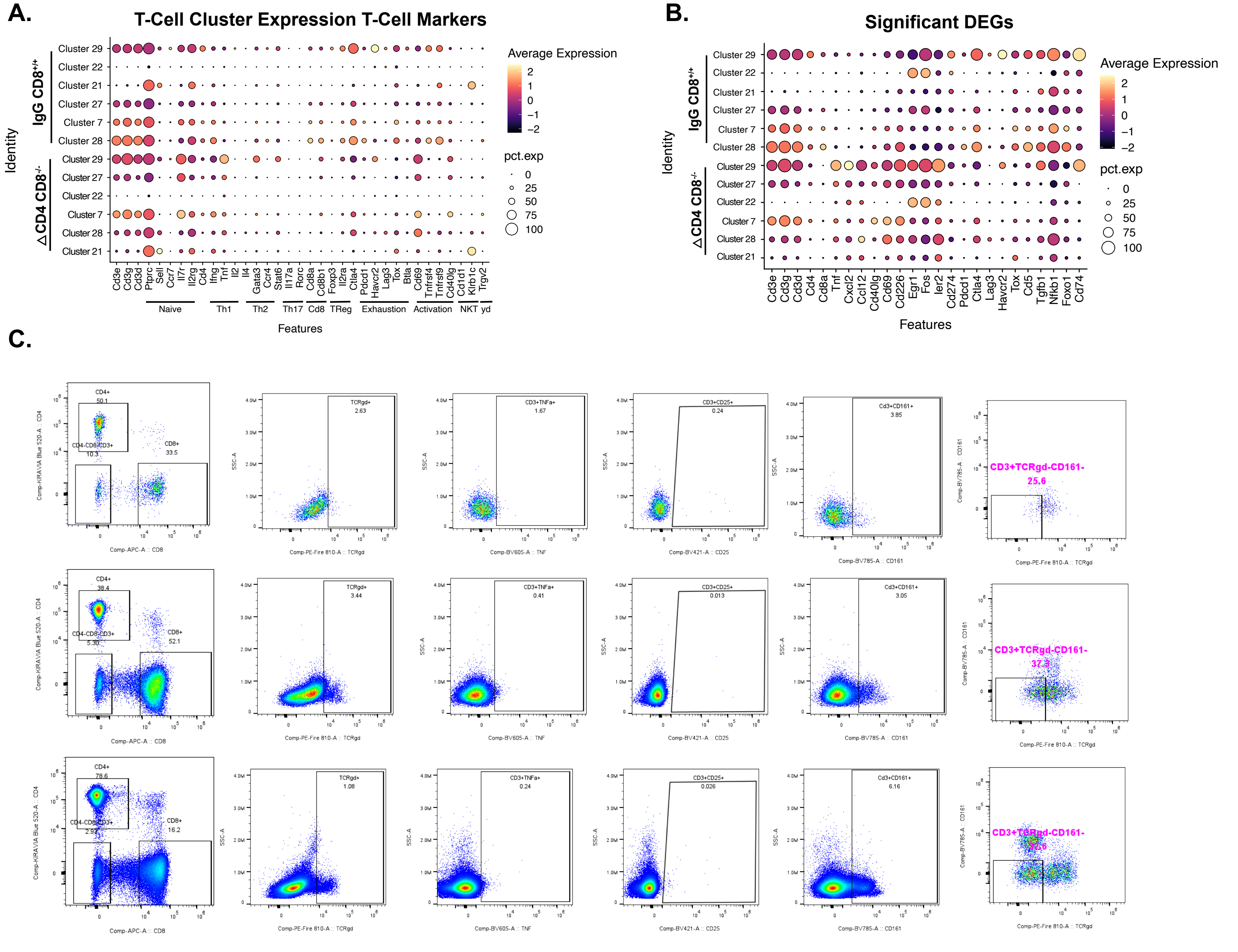
A.
T-Cell Cluster Expression T-Cell Markers
IgG CD8+/+
ΔCD4 CD8-/-
Identity
Cluster 29
Cluster 22
Cluster 21
Cluster 27
Cluster 7
Cluster 28
Average Expression
pct.exp
Cd3e Cd3g Cd3d Ptprc Sell Ccr7 Il7r Il2rg Cd4 Ifng Tnf Il2 Il4 Gata3 Ccr4 Stat6 Il17a Rorc Cd8a Cd8b1 Foxp3 Il2ra Ctla4 Pdcd1 Havcr2 Lag3 Tox Btla Cd69 Tnfrsf4 Tnfrsf9 Cd40lg Cd1d1 Klrb1c Trgv2
Naive
Th1
Th2
Th17
Cd8
TReg
Exhaustion
Activation
NKT
yd
Features
B.
Significant DEGs
Cd3e Cd3g Cd3d Cd4 Cd8a Tnf Cxcl2 Ccl12 Cd40lg Cd69 Cd226 Egr1 Fos Ier2 Cd274 Pdcd1 Ctla4 Lag3 Havcr2 Tox Cd5 Tgfb1 Nfkb1 Foxo1 Cd74
C.
CD4+
CD8+
CD4-CD8-CD3+
TCRgd+
CD3+TNFa+
CD3+CD25+
CD3+CD161+
CD3+TCRgd-CD161-
Comp-APC-A :: CD8
Comp-KIRAVIA Blue 520-A :: CD4
Comp-PE-Fire 810-A :: TCRgd
Comp-BV605-A :: TNF
Comp-BV421-A :: CD25
Comp-BV785-A :: CD161
SSC-A
