## Supplemental Figure 3 for "CD3⁺CD4⁻CD8⁻ (Double-Negative) T-cells Provide Inflammatory Signals That Influence Myeloid Cells and Enhance Survival in Glioblastoma"

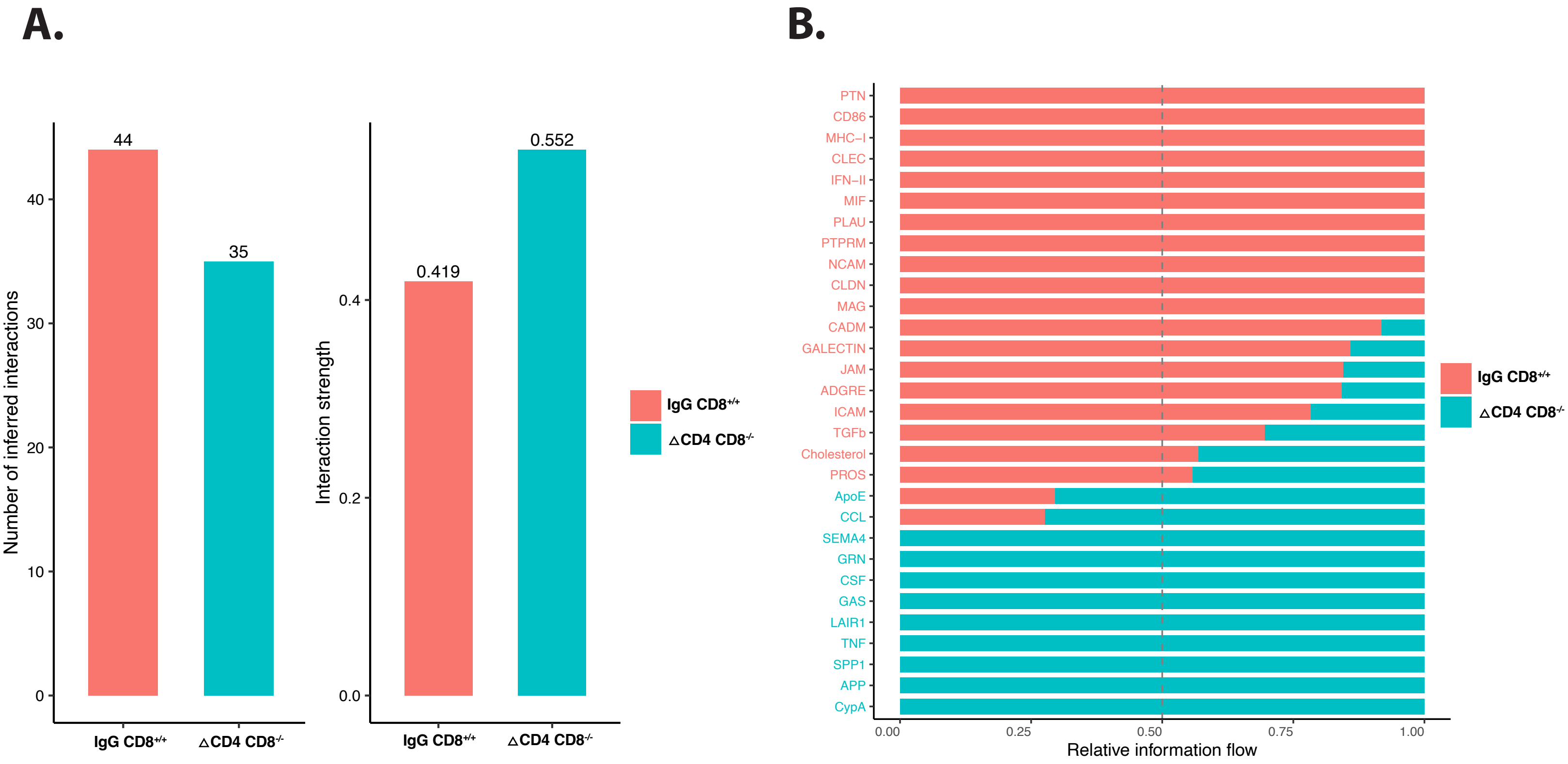

A.
44
35
40
30
20
10
0
Number of inferred interactions
IgG CD8+/+
△CD4 CD8-/-
0.552
0.419
0.4
0.2
0.0
Interaction strength
IgG CD8+/+
△CD4 CD8-/-
IgG CD8+/+
△CD4 CD8-/-
B.
PTN
CD86
MHC−I
CLEC
IFN−II
MIF
PLAU
PTPRM
NCAM
CLDN
MAG
CADM
GALECTIN
JAM
ADGRE
ICAM
TGFb
Cholesterol
PROS
ApoE
CCL
SEMA4
GRN
CSF
GAS
LAIR1
TNF
SPP1
APP
CypA
0.00
0.25
0.50
0.75
1.00
Relative information flow
IgG CD8+/+
△CD4 CD8-/-
