## Supplemental Figure 5 for "CD3⁺CD4⁻CD8⁻ (Double-Negative) T-cells Provide Inflammatory Signals That Influence Myeloid Cells and Enhance Survival in Glioblastoma"

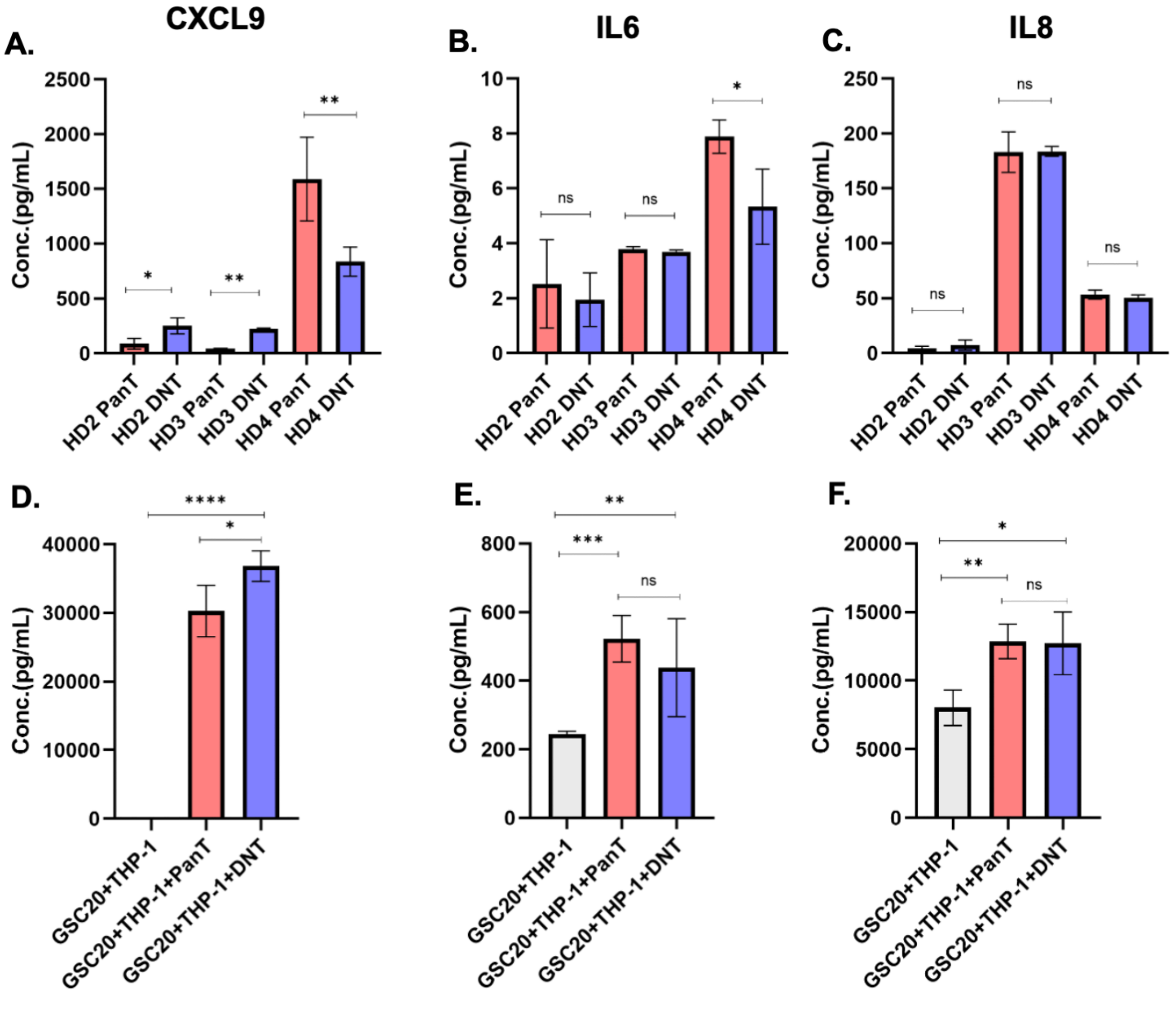
A.
CXCL9
Conc.(pg/mL)
2500
2000
1500
1000
500
0
*
**
**
HD2 PanT
HD2 DNT
HD3 PanT
HD3 DNT
HD4 PanT
HD4 DNT
B.
IL6
Conc.(pg/mL)
10
8
6
4
2
0
ns
ns
*
HD2 PanT
HD2 DNT
HD3 PanT
HD3 DNT
HD4 PanT
HD4 DNT
C.
IL8
Conc.(pg/mL)
250
200
150
100
50
0
ns
ns
ns
HD2 PanT
HD2 DNT
HD3 PanT
HD3 DNT
HD4 PanT
HD4 DNT
D.
Conc.(pg/mL)
40000
30000
20000
10000
0
****
*
GSC20+THP-1
GSC20+THP-1+PanT
GSC20+THP-1+DNT
E.
Conc.(pg/mL)
800
600
400
200
0
**
***
ns
GSC20+THP-1
GSC20+THP-1+PanT
GSC20+THP-1+DNT
F.
Conc.(pg/mL)
20000
15000
10000
5000
0
*
**
ns
GSC20+THP-1
GSC20+THP-1+PanT
GSC20+THP-1+DNT
