## Supplemental Figure 6 for "CD3⁺CD4⁻CD8⁻ (Double-Negative) T-cells Provide Inflammatory Signals That Influence Myeloid Cells and Enhance Survival in Glioblastoma"

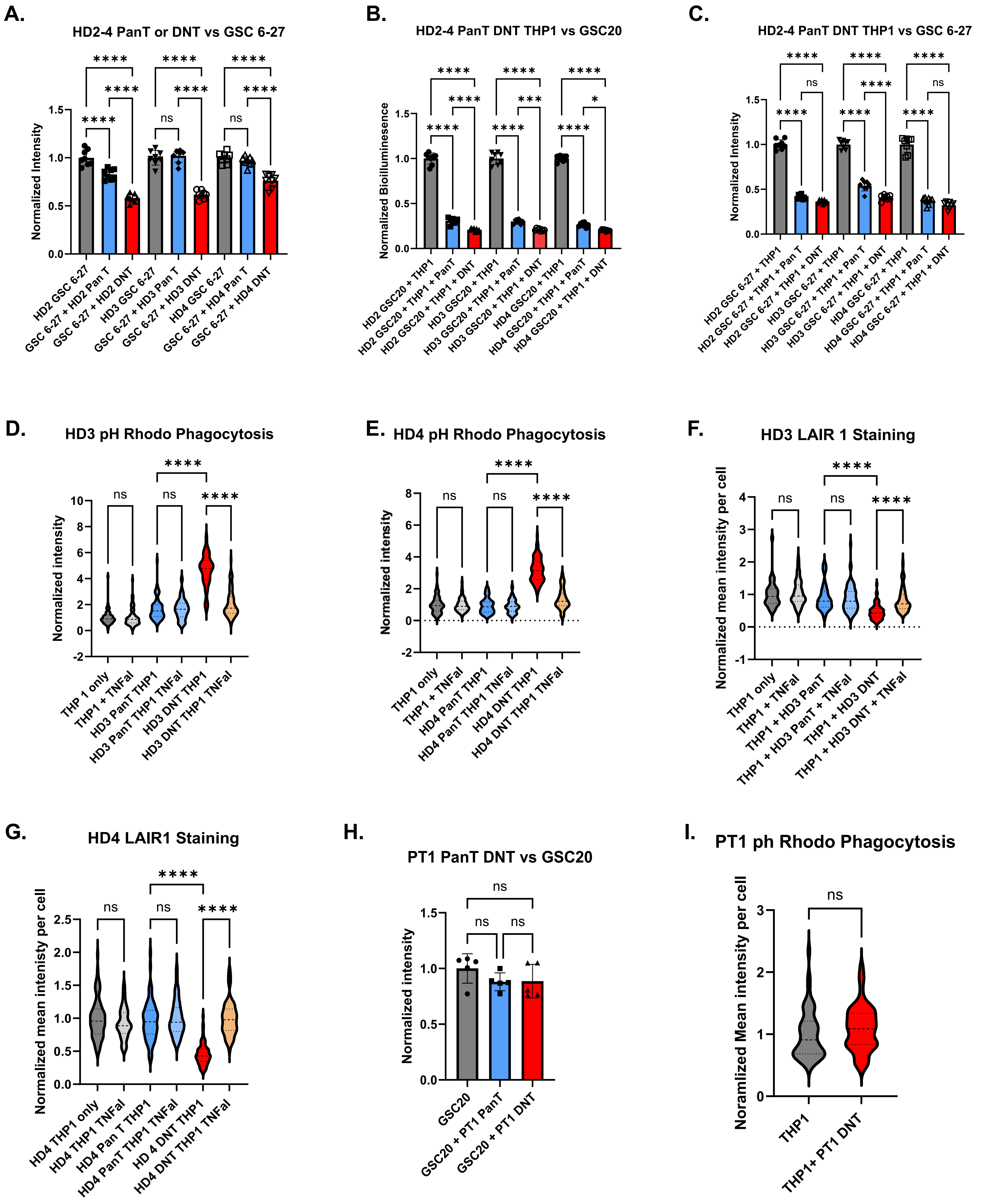
A.
HD2-4 PanT or DNT vs GSC 6-27
Normalized Intensity
HD2 GSC 6-27
GSC 6-27 + HD2 Pan T
GSC 6-27 + HD2 DNT
HD3 GSC 6-27
GSC 6-27 + HD3 Pan T
GSC 6-27 + HD3 DNT
HD4 GSC 6-27
GSC 6-27 + HD4 Pan T
GSC 6-27 + HD4 DNT
B.
HD2-4 PanT DNT THP1 vs GSC20
Normalized Bioilluminesence
HD2 GSC20 + THP1
HD2 GSC20 + THP1 + PanT
HD2 GSC20 + THP1 + DNT
HD3 GSC20 + THP1
HD3 GSC20 + THP1 + PanT
HD3 GSC20 + THP1 + DNT
HD4 GSC20 + THP1
HD4 GSC20 + THP1 + PanT
HD4 GSC20 + THP1 + DNT
C.
HD2-4 PanT DNT THP1 vs GSC 6-27
Normalized Intensity
HD2 GSC 6-27 + THP1
HD2 GSC 6-27 + THP1 + Pan T
HD2 GSC 6-27 + THP1 + DNT
HD3 GSC 6-27 + THP1
HD3 GSC 6-27 + THP1 + Pan T
HD3 GSC 6-27 + THP1 + DNT
HD4 GSC 6-27 + THP1
HD4 GSC 6-27 + THP1 + Pan T
HD4 GSC 6-27 + THP1 + DNT
D.
HD3 pH Rhodo Phagocytosis
Normalized intensity
THP 1 only
THP1 + TNFal
HD3 PanT THP1
HD3 PanT THP1 TNFal
HD3 DNT THP1
HD3 DNT THP1 TNFal
E.
HD4 pH Rhodo Phagocytosis
Normalized intensity
THP1 only
THP1 + TNFal
HD4 PanT THP1
HD4 PanT THP1 TNFal
HD4 DNT THP1
HD4 DNT THP1 TNFal
F.
HD3 LAIR 1 Staining
Normalized mean intensity per cell
THP1 only
THP1 + TNFal
THP1 + HD3 PanT
THP1 + HD3 PanT + TNFal
THP1 + HD3 DNT
THP1 + HD3 DNT + TNFal
G.
HD4 LAIR1 Staining
Normalized mean intenisty per cell
HD4 THP1 only
HD4 THP1 TNFal
HD4 Pan T THP1
HD4 PanT THP1 TNFal
HD 4 DNT THP1
HD4 DNT THP1 TNFal
H.
PT1 PanT DNT vs GSC20
Normalized intensity
GSC20
GSC20 + PT1 PanT
GSC20 + PT1 DNT
I.
PT1 ph Rhodo Phagocytosis
Noramlized Mean intensity per cell
THP1
THP1+ PT1 DNT
